# Accurate Detection of RNA Stem-Loops in Structurome Data Reveals Widespread Association with Protein Binding Sites

**DOI:** 10.1101/2021.04.28.441809

**Authors:** Pierce Radecki, Rahul Uppuluri, Kaustubh Deshpande, Sharon Aviran

## Abstract

RNA molecules are known to fold into specific structures which often play a central role in their functions and regulation. *In silico* folding of RNA transcripts, especially when assisted with structure profiling (SP) data, is capable of accurately elucidating relevant structural conformations. However, such methods scale poorly to the swaths of SP data generated by transcriptome-wide experiments, which are becoming more commonplace and advancing our understanding of RNA structure and its regulation at global and local levels. This has created a need for tools capable of rapidly deriving structural assessments from SP data in a scalable manner. One such tool we previously introduced that aims to process such data is *patteRNA*, a statistical learning algorithm capable of rapidly mining big SP datasets for structural elements. Here, we present a reformulation of *patteRNA*’s pattern recognition scheme that sees significantly improved precision without major compromises to computational overhead. Specifically, we developed a data-driven logistic classifier which interprets *patteRNA*’s statistical characterizations of SP data in addition to local sequence properties as measured with a nearest neighbor thermodynamic model. Application of the classifier to human structurome data reveals a marked association between detected stem-loops and RNA binding protein (RBP) footprints. The results of our application demonstrate that upwards of 30% of RBP footprints occur within loops of stable stem-loop elements. Overall, our work arrives at a rapid and accurate method for automatically detecting families of RNA structure motifs and demonstrates the functional relevance of identifying them transcriptome-wide.

## INTRODUCTION

Beyond serving as a carrier of genetic information, RNA plays key mechanistic roles in diverse cellular processes. These functions are regularly attributed to the molecule’s ability to fold into specific structures (1–7). Driven by its flexible backbone and the complementarity of nucleotide bases comprising it, the structures of RNA are intricate and dynamic (1, 8). Although high-quality structure models of RNA transcripts are important in understanding their function and dysfunction, accurate determination of structures, especially *in vivo*, is challenging. High-resolution structure models can be obtained with experimental measurements from X-ray crystallography (9), nuclear magnetic resonance (10), and cryo-EM (3, 11), yet these methods are low-throughput and incapable of measuring structures in living cells. Comparative sequence analyses can also glean structural information from sequence homologies, but this process depends on a sufficiently large and suitably divergent set of related sequences, which limits the scope of their application (12–14). The advent of nearest-neighbor thermodynamic models (NNTM) combined with efficient energy minimization algorithms were a critical step in increasing the throughput of structural predictions by enabling computational folding based on nucleotide sequences (15, 16). Despite their popularity, however, the accuracy of predictions is generally poor, especially when applied *in vivo* or to long transcripts (17). Structure profiling (SP) experiments have emerged as a practical and high-throughput approach to measuring the structure of RNA molecules (18, 19). Although these methods are diverse, they help inform structure models by providing nucleotide-level measurements of conformational characteristics. Importantly, they are broadly applicable (e.g., viable *in vivo* or in other conditions), and, with the advent of next-generation sequencing, are scalable to entire transcriptomes.

SP experiments follow common principles (20). Briefly, they expose RNA to chemical reagents or enzymes that react with parts of the molecule in a structure-dependent manner; for example, when using common acylation reagents, single stranded nucleotides react more strongly than double stranded regions. This reaction induces the formation of adducts or cleavages, which can then be detected during sequencing as either truncations or mutations in reverse-transcribed cDNA fragments (18–23). The rate of truncation or mutation at each nucleotide is then quantified and converted into a measure called reactivity that summarizes the nucleotide’s structural context; the reactivities across a transcript are termed its reactivity profile (20, 24). The incorporation of these data in NNTM-based folding algorithms was shown to greatly improve their accuracy (25, 26). In this regard, SP data have traditionally served to supplement the thermodynamic models by providing direct information on the measured conformation. That said, SP experiments have scaled massively, enabling the profiling of an entire transcriptome, termed the structurome. NNTM-based folding, however, is a computationally intensive process that scales as *O*(*L*^3^) with the length of an RNA in most applications. For transcriptomes, which contain many tens of thousands of transcripts— each of which may be thousands of nucleotides long—the computational cost associated with folding has begun to inhibit comprehensive NNTM-based analyses of structurome data. This has warranted the development of methods designed to accommodate the growing scale of SP data in deciphering the extensive regulatory functions of mRNA structures. Such methods are useful when seeking to inspect and quantify biologically relevant changes in the structurome—for instance, to detect structural changes between different cellular conditions (27–30), inspect the structural context of relevant regions, such as splice sites, miRNA targets, or alternative polyadenylation sites (31–33), profile the degree of structure across different types of mRNA (33), or explore the interplay between the structurome and RNA-protein interactome (34).

We previously introduced *patteRNA* as a method to address this need for scalable analysis of SP data (35). Rather than perform complete RNA folding, it was developed to rapidly mine local structure elements from reactivity profiles via an unsupervised, versatile, and NNTM-free approach. In short, the method couples a statistical reactivity model—e.g., a Gaussian mixture model (GMM) or discretized observation model (DOM)—to a Hidden Markov model of structure (35–37). A parameterized model subsequently enables rapid quantitative scanning for locations that are likely to harbor a specific structure element. Versatility is a key characteristic; namely, it leverages an unsupervised training step to learn the properties of any type of SP dataset (i.e., to parameterize the reactivity-structure model) before mining it. This is crucial for the automated and adaptive handling of data from diverse SP experiments that consequently have disparate statistical properties. Currently, standard practice is to use methods tailored for specific reactivity distributions (i.e., *in vitro* SHAPE) characteristic of highly structured non-coding RNAs. Such workflows do not capture the full diversity of probes, conditions, and SP pipelines, rendering them suboptimal, especially for mRNA-centric *in vivo* structurome studies (38). Moreover, the NNTM-free nature of *patteRNA* helps it scale to the structurome level and also confers flexibility to rapidly mine complex structural elements such as pseudoknots or self-contained tertiary interactions without any significant increase in computational complexity (35). In short, any target that can be defined via a local reactivity pattern can be quickly mined.

By scanning reactivity profiles alone, *patteRNA* was able to achieve reasonable accuracy when mining canonical motifs, such as hairpins/stem-loops (37). However, there was room for improvement via integration of NNTM-derived sequence information, which we believed could likely assist in situations where SP data is inconclusive. However, effective integration of NNTM with the statistical framework underpinning our approach is itself non-trivial. This is because we sought to not only improve performance, but also to maintain speed and versatility. To address this, we took a data-driven approach in which a large set of reference structures guided the construction of an integrative scoring classifier which considers statistical characterization of SP data in additional to local thermodynamics. This is a deviation from the unsupervised nature of our approach; nevertheless, we ensured that the classifier maintains the method’s automated adaptability in analyzing any type of SP dataset. The impact of including thermodynamics on the method’s efficiency was also carefully considered, as we sought to maintain a balance between improvements to prediction quality and the increased computational overhead triggered by thermodynamic modeling.

Our results describe the development of a data-driven logistic regression classifier to better identify the locations of target structural elements. It considers the thermodynamic properties of local regions in addition to reactivity profiles when making predictions, which strongly improves precision, especially for shorter structure motifs. The classifier is suitable for all types of canonical local structure motifs and maintains the versatility of *patteRNA* in handling diverse types of SP data. In this process, we also create a large-scale set of RNAs with known structures from RNA STRAND (39) and use it in conjunction with data simulations to extensively train and validate our approach. Although underpinned by simulated data, we find this resource to be more effective at training data-driven classifiers than smaller sets of real data and believe it can serve as a useful resource for machine learning method development. Moreover, we apply the classifier to an integrative transcriptomic dataset on human cells that quantifies both structure and RNA binding protein (RBP) interactions (34). We demonstrate that stable stem-loops are almost always associated with evidence of RBP binding, and that this association exists across a diverse set of stem-loop configurations. In the context of the latest RBP studies, our results expand on previous observations of the RNA-protein interactome and refine our understanding of the roles played specifically by stem-loops. This also highlights the power of stem-loop profiling, a task for which alternative specialized tools are currently lacking. Overall, our work provides a major improvement to *patteRNA* while simultaneously enhancing our understanding of the functional roles canonical structure elements play in RNA-protein interactions.

## MATERIALS AND METHODS

### patteRNA Overview

*patteRNA* works in two phases: training and scoring. The training phase involves the utilization of an unsupervised Expectation-Maximization (EM) scheme coupled to a Hidden Markov Model (HMM) to estimate the reactivity distributions for unpaired and paired states, respectively. With these distributions in hand, *patteRNA* searches for a target motif in SP data as previously described (35, 36). Briefly, all subsequences (referred to as sites) which satisfy the sequence constraints underlying the base pairing arrangement of the target motif are considered. These sites are then each assigned a score, which quantifies the overall consistency of the reactivity data within the site with the pairing state sequence of the target (a higher score indicates a better agreement between the reactivity profile and target motif). Scores are further processed into *c*-scores via a normalization scheme based on an estimated distribution of scores associated with null sites (sites that do not harbor the target motif). For details on the core *patteRNA* algorithm, see (35) and (37); for details on score normalization, see (36).

All applications of *patteRNA* in this study used default hyperparameters unless otherwise noted. When mining hairpins, the “--hairpin” flag was used, which searches for all hairpins/stem-loops with stem length between 4 and 15 nt and loop length between 3 and 10 nt. This representative collection of motifs is referred to as regular hairpins or regular stem-loops throughout our work. When mining loops, the “--loops” flag was used, which searches for runs of unpaired nucleotides length 3 to 10 nt flanked by one base pair.

### The Weeks Set

The Weeks set is a dataset of 22 diverse RNA transcripts (totaling 11,070 nt) with high-quality SHAPE data and known reference structures. We use the Weeks set in this study as a reference set to benchmark the performance of *patteRNA*’s analyses and related methods on real data. This dataset was initially introduced in (35) and contains reactivity data from (25, 40, 41), see **Supplementary Table S1** for further details on the RNA molecules in the Weeks set.

### Classifier Training Data

In order to construct a larger set of reference data by which to develop a scoring classifier, we compiled all RNA secondary structures from RNA STRAND (4,666 transcripts). Due to the presence of highly similar sequences within the data, we used CD-HIT-EST (42) to remove redundant sequences at an 80% similarity threshold, yielding 1,191 final transcripts (totaling 706,306 nt). In order to utilize these secondary structures for *patteRNA*-related analyses, we generated artificial SP data for the transcripts according to a three-state reactivity model (0: unpaired, 1: paired, 2: helix-end) with associated state reactivity distributions devised in (43), which we refer to as the Heitsch distributions. The distributions are defined as follows; unpaired states: exponential distribution with *λ* = 1.468, paired state: generalized extreme value distribution with *μ* = 0.04, *σ* = 0.040, *ξ* = −0.763, helix-end state: generalized extreme value distribution with *μ* = 0.09, *σ* = 0.114, *ξ* = −0.821. Five replicates of SP data were produced. The Python module SciPy was used to sample reactivities from the corresponding distributions. The scripts used to sample reactivities for STRAND transcripts in addition to the STRAND data itself (including the sampled reactivities used in this work) are available at http://doi.org/10.5281/zenodo.4667909, reference number (44). To assist in verification and benchmarking of classifiers, additional datasets were also generated by resampling (with replacement) the empirical reactivity distributions observed in the Weeks set.

### Feature Generation

Several features were investigated insofar as their potential to provide additional information on the presence of target motifs during scoring. After preliminary investigations, we focused on the following features, in addition to the *patteRNA c*-score: cross-entropy loss (CEL) between *patteRNA* posteriors and the target state sequence, Gini coefficient of SHAPE data in a site, the local minimum free energy (LMFE), the local constrained minimum free energy (LCMFE; the local MFE with the target motif enforced as a folding constraint), and the motif energy loss (MEL; the difference between LMFE and LCMFE). Cross-entropy loss was computed as

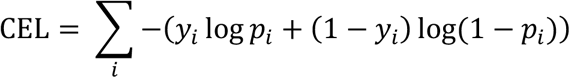

where *y*_*i*_ is the pairing state of the target motif (e.g., *y*_*i*_ = 0 for unpaired states and *y*_*i*_ = 1 for paired) and *p*_*i*_ is the posterior pairing probability at nucleotide *i* of a scored site. The Gini coefficient was computed as

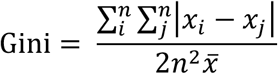

where *x*_*i*_ is the reactivity at nucleotide *i* of a site and *n* is the length of the target motif. The remaining three features (LMFE, LCMFE, and MEL) all depend on the thermodynamic model employed in RNA structure prediction and were computed using the ViennaRNA package (version 2.4.17) Python interface using a local window extending 40 nt in both directions from the boundaries of the target site (*c* = 40).

### Feature Selection

To identify the set of features which best predict the presence of a target motif, we used a factorial-like approach to test various combinations of features and their potential scoring efficacy. To do this, we used the scoring feature set generated from the Weeks set hairpins. We used the *c*-score as a base feature in all experiments while iterating through pairwise combinations of the other features on top of it (see **Figure 1**). Specifically, we tested all of the 2-feature approaches underpinned by the *c*-score and one of the other features. We then tested all of the 3-feature approaches underpinned by the *c*-score and all of the pairwise combinations of the other features. To quantify scoring efficacy, a logistic classifier was trained on the feature combinations and the average precision of its predicted motif probabilities was used to assess scoring potential.

**Figure 1:**
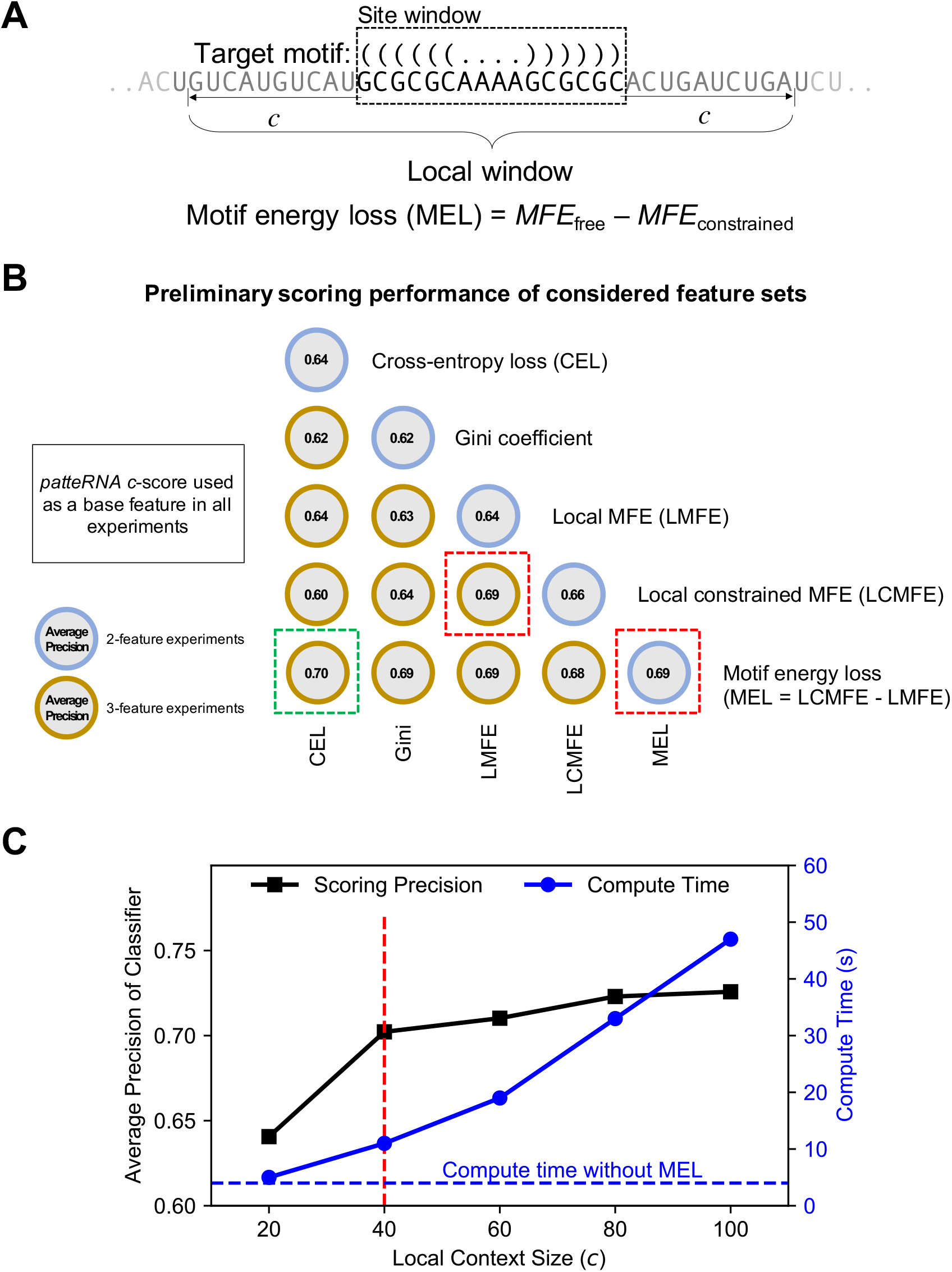
Auxiliary feature development for assisting in structure motif mining from SP datasets. (A) Illustration of the local window considered when computing thermodynamics-based features for scoring, such as motif energy loss (MEL). The considered window extends a distance, *c* (the local context size), from the boundaries of the scored site. (B) Preliminary scoring performance of considered feature sets. A combinatorial approach was used to test the performance of feature combinations. Features were benchmarked by using them to train a logistic classifier and then computing their average precision on a hairpin test set across five replicates; mean average precision is shown. (C) Determination of a suitable context size to use for MEL computations in *patteRNA*. Shown is the scoring precision when using a logistic classifier with *c*-score, cross-entropy loss (CEL), and MEL across various context sizes. Also shown is the measured runtime at each context length. Highlighted in red is the chosen default context size (40 nt), which strikes a balance between scoring precision and computational overhead relative to *patteRNA*’s original speed on the utilized data.

### Classifier Selection

After converging on a 3-feature set of *c*-score, CEL, and MEL, we explored the capacity of various binary classifiers to precisely identify true positive sites from these features as well as their ability to generalize to other datasets and target motifs beyond hairpins. After a preliminary analysis of an initial collection of standard classifiers, we explored in more detail logistic binary classification (LBC; “LogisticRegression” object in Scikit-learn), random forest classification (RFC, “RandomForestClassifier” object in Scikit-learn), linear discriminant analysis (LDA, “LinearDiscriminantAnalysis” object in Scikit-learn), and quadratic discriminant analysis (QDA, “QuadraticDiscriminantAnalysis” object in Scikit-learn). In all cases, default parameterizations were used as provided by Scikit-learn.

The set of hairpins mined from the STRAND dataset was used to train each classifier, and the average precision of their trained predictions was computed. Trained classifiers were then tested against the generated feature set for Weeks set hairpins and for Weeks set loops. Lastly, the classifiers were verified against feature sets obtained when mining 5 resampled replicates of the empirical data in Weeks set (for hairpins and for loops) and when mining 5 replicates of simulated STRAND sets (for hairpins and for loops). In all cases, performance was assessed by the average precision of the classifier.

### Final Scoring Classifier Training and Selection

To train the definitive classifier used in *patteRNA*, we utilized the 5 replicates of Heitsch-simulated reactivity data for the pruned STRAND transcripts and used each replicate to generate a scoring feature set at the sites scored when mining for hairpins. These scoring feature sets were each used to train an associated logistic classifier – i.e., 5 distinct classifiers were trained simultaneously, 1 for each replicate of simulated data. Each resulting classifier was then used to process the other four feature sets (i.e., the other simulations not used to train that classifier) as well as the feature set associated with the empirical Weeks set data. The overall performance of the classifiers was assessed as the sum of the performance on the other 4 STRAND replicates (computed as the mean average precision for hairpins across the 4 replicates) and the performance on the Weeks set (average precision for hairpin mining). The classifier with the greatest total performance via this assessment was selected as the final model to utilize for distribution in the *patteRNA* method.

### Performance Benchmarks and Verification

The accuracy of *patteRNA* and tested binary classifiers was primarily assessed via the area-under-the-curve of the precision-recall (PR) curve, referred to as the average precision (AP) of the classifier. Precision-recall curves were computed by varying a theoretical score threshold between positives and negatives, then computing the true-positive rate (recall) and precision (PPV) at each threshold. Sites were deemed true positives if all base pairs in the target motif are also present in the corresponding location of the reference structure. The Scikit-learn Python module was utilized to perform these computations. Scripts that perform this quantification (and others, including the receiver-operating characteristic) are available in (37).

### Partition Function Analysis

We benchmarked the performance of partition function approaches to detect hairpins in the Weeks set by using the “RNAsubopt” command from ViennaRNA to generate 1000 structures for each transcript in the Weeks set, using that transcript’s SHAPE data as soft constraints (“RNAsubopt -p 1000 --shape ${SHAPE_FILE} < ${SEQUENCE}”). For each hairpin in the generated structural ensemble, a “score” was assigned as the fraction of structures in the structural ensemble which contain the base pairs comprising that hairpin. Predicted hairpins and their scores were organized into a single list which was then processed into a precision-recall curve as was done for *patteRNA*’s predicted hairpins.

### Analysis of Structurome and RBP Binding Data

We used *patteRNA* with the latest logistic scoring classifier described above to mine hairpins in the *in vitro* and *in vivo* icSHAPE data from K562 and HepG2 cells published by Corley et al. (34). *patteRNA* was trained on each dataset/condition independently (e.g., K562 *in vitro* icSHAPE, K562 *in vivo* icSHAPE, HepG2 *in vitro* icSHAPE, etc.) and then used to mine them for hairpins (referred to in this analysis as stem-loops) using the “-hairpins” flag and default hyperparameters. We then cross-referenced the locations of high-scoring stem-loops with the fSHAPE profiles (interpreted as RBP binding signal) obtained by Corley et al. on the same cell lines.

To combine and visualize the fSHAPE profiles from searched sites which differ in terms of their stem and loop lengths, we utilized an interpolation scheme to bring fSHAPE profiles to a common length basis. fSHAPE profiles from the left and right sides of the stem (which vary from 4 to 15 nt in length) were each processed to a length of 10, respectively. fSHAPE profiles from the loop regions were processed to a length of 6. This processing was achieved by linearly interpolating the fSHAPE profiles to a number of equally spaced points (e.g., 10 points for stems and 6 points for loops). For example, a stem of length 6 nt would be linearly interpolated to the local coordinates (1, 1.56, 2.11, 2.67, 3.22, 3.78, 4.33, 4.89, 5.44, 6), where 1 and 6 denote beginning and end of the fSHAPE profile along the stem, respectively.

Motif scores from both conditions were then combined and used to train a perceptron classifier processing condition-wise paired scores from the LBC into a predictor of strong RBP binding signal in the loop (defined as sites where fSHAPE > 2 in the stem-loop). Only sites that received a valid score in both conditions were considered in this analysis. The multi-layer perceptron (MLP) classifier object (MLPClassifier) from Scikit-learn was utilized to construct and train the classifier; the default model parameterization was used, which is defined by a single hidden layer of 100 nodes with ReLU activation following the Adam optimization algorithm (45). Cross-validation during perceptron training was achieved by randomly setting aside 20% of the samples and using them to terminate training when convergence was observed (this behavior was defined with the hyperparameters “validation_fraction=0.2” and “early_stopping=True” when calling MLPClassifier). Note that the purpose of the perceptron in this case is, essentially, to fit the joint distribution of condition-wide scores as an indicator of high loop fSHAPE. We found that this non-linear relationship of scores between conditions was best captured by a simple perceptron instead of simpler linear models like logistic regression and LDA.

*patteRNA* was also used to mine the icSHAPE data for stem-loops with bulges, which we defined as stem-loops with stem length between 5 and 15 nt with one bulge (of 1-2nt) on either side of the stem. Locations of high scoring motifs were then cross-referenced against the locations of high fSHAPE, as was done for the hairpin search.

The proportion of strong RBP binding signals (defined as fSHAPE > 2) which can be explained as occurring within the loop of a detected stem-loop was quantified. In this quantification, only fSHAPE observations which coincide with valid reactivity data were included. In other words, fSHAPE data at locations lacking reactivity information was omitted, as such regions are not processed by *patteRNA* when scoring. Three score thresholds for calling detected stem-loops were used: 0.9, 0.7, and 0.5.

Stem-loop detection rates were computing by counting the number of unique stem-loops detected within each logical mRNA region (5’ UTR, CDS, 3’ UTR, with non-coding RNAs treated as their own region) and dividing totaled counts by the number of reactivity observations within each region (to account for the fact that *patteRNA* only considers sites with SP data). Results were multiplied by 1000 to arrive at more interpretable rates in units of stem-loops per 1000 nucleotides.

## RESULTS

### patteRNA Overview

The overarching objective of *patteRNA* is to accurately mine structure elements from SP data in an automated fashion. To do this, *patteRNA* follows a two-step process (see **Supplementary Figure S1**). The first step is the training phase, during which reactivities are utilized to iteratively optimize a joint reactivity-structure statistical model (e.g., a GMM-HMM (35, 46) or a DOM-HMM (37)). This results in an estimate of the distributions of reactivities associated with paired and unpaired nucleotides, respectively, as well as transition probabilities between paired and unpaired nucleotides. Training is unsupervised and capable of accommodating diverse data types; see (35) for a complete description of the mathematical framework.

Once the data properties have been learned, *patteRNA* mines for structural motifs in the data via a scoring step. Scoring requires the description of a specific secondary structure motif (or collection of motifs) which defines the target of *patteRNA*’s pattern recognition scheme. Typically, the user provides this motif in dot-bracket format, but *patteRNA* also has built-in routines to automatically mine some canonical motifs. For instance, *patteRNA* can automatically mine a representative set of hairpins (referred to as regular hairpins or regular stem-loops; defined as stem-loops with stem length between 4 and 15 nt, and loop length between 3 and 10 nt) via the “--hairpins” flag (37). When mining a particular structural element, only loci in the provided transcripts which satisfy the sequence constraints necessary for the target’s secondary structure (via Watson-Crick and Wobble base pairs) are considered during scoring; these loci are henceforth referred to as “sites.” *patteRNA* scores sites by computing the log ratio of joint probabilities between the target’s pairing sequence and its inverse. By default, scores are further processed into *c*-scores (comparative scores) which are a statistically-normalized measure computed by considering the significance of a score in the context of a null-score distribution constructed for each target (36). Intuitively speaking, *c*-scores are simply the −log_10_ of a *p*-value, facilitating comparative analysis of scores from different target searches. Higher scores indicate a higher likelihood of the target motif, with a *c*-score of 2 (corresponding to a *p*-value of 0.01) generally interpretable as a strong signal.

In addition to scoring, *patteRNA* can also use a trained statistical model to compute posterior pairing probabilities (i.e., for each nucleotide, the probability that it is paired or unpaired), Viterbi paths (the most likely sequence of pairing states for each transcript), and hairpin-derived structure level (HDSL) profiles (a nucleotide-wise measure of local structuredness (37)).

### Supervised Context-Aware Scoring

*patteRNA* was developed as an NNTM-free method. It inspects and quantifies patterns in reactivity profiles to identify sites consistent with the presence of a sought structure motif. Sequence information is only taken into account when determining whether sites are compatible with a target motif, i.e., satisfying the sequence constraints associated with base pairs in the target. This approach has facilitated the algorithm’s rapid speed when mining transcriptomic data; however, information encoded in the sequence has the potential to improve its predictions. Here, we aimed to improve *patteRNA’s* accuracy by including an assessment of information in nucleotide sequences (e.g., NNTM-based quantifications of sequence energetics). We also explored the use of additional SP data-related metrics in improving performance.

The integration of NNTM-based predictions with a statistical metric like the *c*-score is non-trivial. Therefore, we pursued the development of a data-driven scoring classifier, through which multiple features from sites would be processed in assessing the likelihood of a motif. This is a departure from the unsupervised nature of *patteRNA*. Despite this departure, we sought to maintain the broad applicability of the method to diverse data types. As such, we focused on features that we believed to generalize well across SP datasets (i.e., are data invariant).

We explored various features in conjunction with the *c*-score to underpin the classifier. Five features emerged as promising candidates and their potential was further explored in a combinatorial set of experiments. The first was the cross-entropy loss (CEL) between the target motif’s pairing sequence and *patteRNA*’s computed posterior pairing probabilities (see Methods). This feature relates closely to the *c*-score but highlights the cumulative disagreement between the data and the motif more explicitly. Specifically, CEL is influenced more strongly by nucleotide-level disagreements (compared to agreements, see **Supplementary Figure S2**) which may otherwise be masked by the *c*-score. As such, we speculated that this metric could assist scoring as it helps discriminate between sites that score moderately well across their entire span and sites that score strongly for some nucleotides but have strong disagreement in others. This is particularly relevant in the context of RNA structure where distinct motifs are often highly similar outside of a small number of decisive nucleotides. For example, when determining the length of a loop within a stem-loop, nucleotides near the end of the stem may inform the precise extent of the loop (e.g., a loop length of 4 nt versus 6 nt). Local disagreement can distinguish between such competing structures.

The second feature was the Gini index of the reactivities at the target site, which is often used in the context of reactivity analysis (20, 47). Gini index has previously been used to assess structural homogeneity. For instance, we expect stable conformations to yield more distinct reactivities between paired and unpaired states (high Gini index) and less stable structures or regions characterized by multiple conformations yield more intermediate values (low Gini index) (48). As such, we speculated that Gini index could serve as a simple proxy for data quality and structural homogeneity in a site, and therefore might assist in informing where a *c*-score is more or less meaningful.

The third, fourth, and fifth features relate to the thermodynamic prediction of the local region’s minimum free energy (MFE) structure. It has been shown that incorporating thermodynamic models with SP data tends to improve predictions (25, 38, 49). Therefore, we utilized their predictions in different ways to potentially assist as features in a scoring classifier. As such, the third feature was the local minimum free energy (LMFE) of the region around a site, where local is defined as the target site window extended in both directions by some distance (e.g., 40 nt; see **Figure 1A**). The fourth feature was the local constrained minimum free energy (LCMFE) of the region around the site, which amounts to folding with the target motif strictly enforced as a hard constraint. We thought that these two metrics, or perhaps their combination, could assist in interpreting the stability of the local region and the motif’s influence on it. We also considered a fifth feature, which was the difference between LMFE and LCMFE, which we termed the motif energy loss (MEL). This measure summarizes the energetic favorability associated with the presence of the motif.

### Feature Selection

To test the scoring potential of various feature combinations, we established a simple train-and-test pipeline for mining hairpins in a reference dataset (the Weeks set, see Methods). Various feature combinations were used to train a logistic classifier whose scoring precision was then quantified (using an 80%/20% test/train split). For each feature combination, this procedure was repeated 5 times. Mean scoring precision on the test sets was then used to assess the scoring potential of that feature combination.

We performed benchmarks in a simple combinatorial manner to investigate which features and feature combinations were most effective. The *c*-score was used as a base feature in our analysis, meaning that it was included in all combinations. The results of our preliminary feature analysis are in **Figure 1B**. In the 2-feature experiments, all candidate features except Gini index yielded a detectable improvement in precision over just using the *c*-score (which achieves an average baseline precision of 0.62), and we found that MEL yielded the strongest enhancement (to an average precision of 0.69, an 11% improvement over baseline). The 3-feature experiments were only able to incrementally improve scoring precision beyond this level. The best 3-feature combination was *c*-score, CEL, and MEL, which yielded an average precision of 0.70. Interestingly, the combination of *c*-score, LMFE and LCMFE yielded an average precision approximately equal to the observed precision with *c*-score and MEL. We also observed that none of the 4, 5, or 6-feature classifiers significantly outperformed the best 3-feature classifier on any of the benchmarks (data not shown), further validating the efficiency of the chosen scheme. We chose this set of features to use as inputs when developing and optimizing the scoring classifier to utilize in *patteRNA*’s scoring pipeline.

To determine an appropriate local context size to use for MEL, we investigated the precision of the selected feature set at regular intervals of local context length from 20 nt to 100 nt (note that the full context size used for folding is 2*c* + *n*, where *c* is the context length and *n* is the motif length). We simultaneously measured the respective compute times. Our results, shown in **Figure 1C**, demonstrate a trade-off between feature quality and compute time as one increases the local context size. We observe that the scoring quality plateaus approximately at *c* = 40, yet the extra compute time (relative to NNTM-free scoring) rapidly grows for longer context lengths. For this reason, we decided to use *c* = 40. We note, however, that larger contexts do provide a slightly better structural interpretation. Thus, although a length of 40 nt is used for the remainder of our work in the manuscript, this parameter may to be tuned by the user when calling *patteRNA*.

### Classifier Selection and Optimization

Having converged on using *c*-score, CEL, and MEL, we devised a more intensive classifier training pipeline and used it to investigate a set of standard binary classifiers for their ability to robustly model these features. Specifically, we examined logistic binary classification (LBC), random forest classification (RFC), linear discriminant analysis (LDA), and quadratic discriminant analysis (QDA).

Our classifier training pipeline was underpinned by the use of RNA STRAND (39). STRAND has 4,666 high-quality secondary structure models spanning a large set of RNA families, including regulatory elements, ribosomal RNA, ribozymes, synthetic RNAs, and more. After removal of highly redundant sequences with CD-HIT-EST (42), 1,191 transcripts remained, providing a much more expansive structural snapshot to use for classifier training than the Weeks set, which comprised 22 transcripts. Importantly, STRAND transcripts do not generally have SP data associated with them. Thus, we utilized simulations to generate artificial data. Reactivities were modeled as (and sampled from) the three-state model (unpaired, stacked, and helix-end) devised in (43).

**Figure 2A** demonstrates the interplay between the Weeks set data and the STRAND transcripts as used in our analysis. In short, we used simulated data on STRAND transcripts for classifier training. The Weeks set was then used to benchmark the performance of classifiers trained from STRAND simulations. We found that using STRAND transcripts (with simulated data) yielded the best results in terms of performance on the Weeks test set benchmarks, even outperforming classifiers trained on the Weeks set directly. Verification sets were also generated by resampling additional replicates of the Weeks set and simulating additional replicates on STRAND (see Methods for details). The overall objective was to identify the best possible motif classifier for the 3 investigated features (**Figure 2B**).

**Figure 2:**
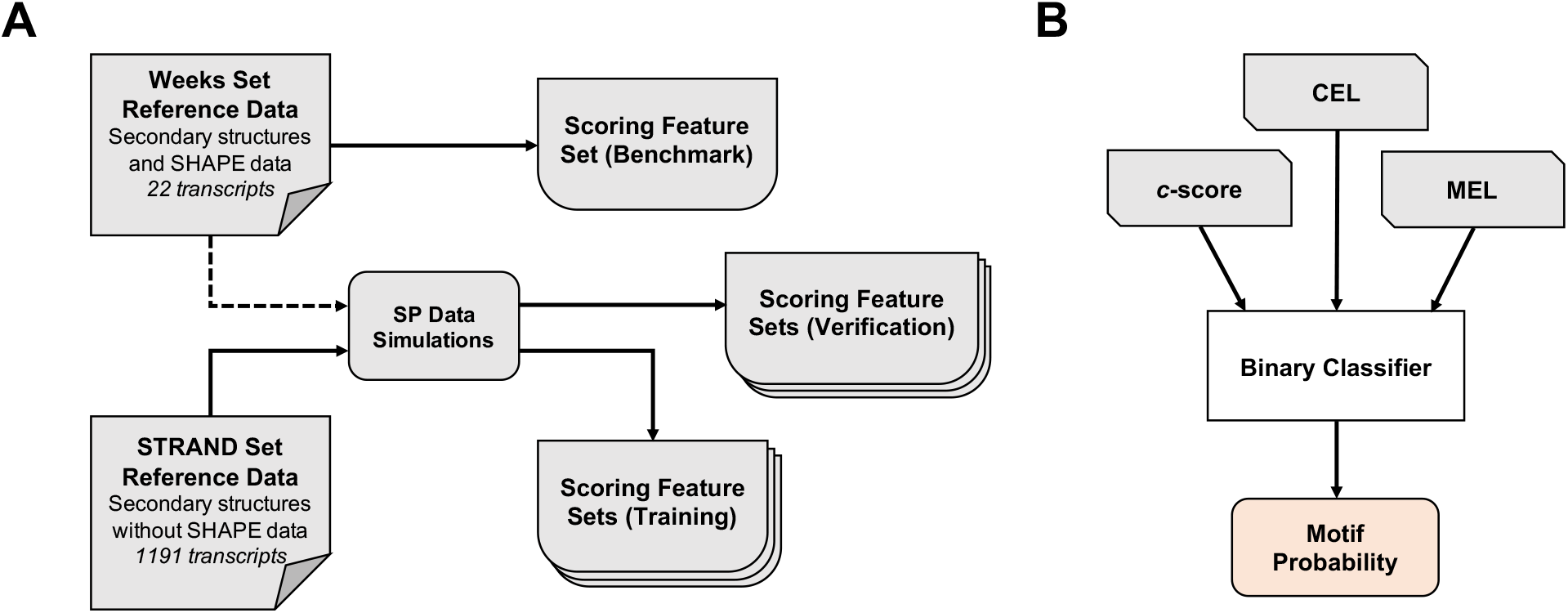
Data processing scheme for feature set generation in training, verifying, and benchmarking a binary motif classifier. (A) Data sources and computational flow for generating features sets used for training, benchmarking, and verification. Transcripts in RNA STRAND were used for classifier training; however, because these transcripts lack SP data, simulations were used to generate artificial reactivities on known secondary structures. The Weeks set was used to benchmark classifiers as it contains RNAs with known structures and high-quality real-world reactivity data. The reactivities in the Weeks set were also resampled to generate additional replicates, which were also used for verification steps in addition to replicated simulations on RNA STRAND. (B) Schematic of the binary classification approach utilized in *patteRNA*. *c*-score, CEL, and MEL were used as the features driving assessments of motif probability.

Our results are presented in **Figure 3**. Overall, we found that the LBC provided the best results in terms of scoring consistency and translatability to verification benchmarks against other data and other motifs. Generally, we observed similar results for LBC, LDA, and QDA—all classifiers strongly improved scoring when compared to *c*-scores on the benchmark and verification sets—yet the LBC slightly exceeded the others’ performance on all tests. We also observed that the LBC was the fastest of the tested classifiers (data not shown). Interestingly, we observed that a random forest classifier was able to achieve remarkable performance on the training set but did not translate effectively to other benchmarks or validations. We presume that the classifier was overfitted due to its parameterization (described by a large number of decision trees); efforts to reduce the size and complexity of the parameterization (e.g., by reducing the number of estimators) were unsuccessful in improving performance beyond what was observed with logistic regression. Moreover, we found the compute time in applying random forest classification to scale poorly in situations where a large number of sites (i.e., more than tens of thousands) were scored.

**Figure 3:**
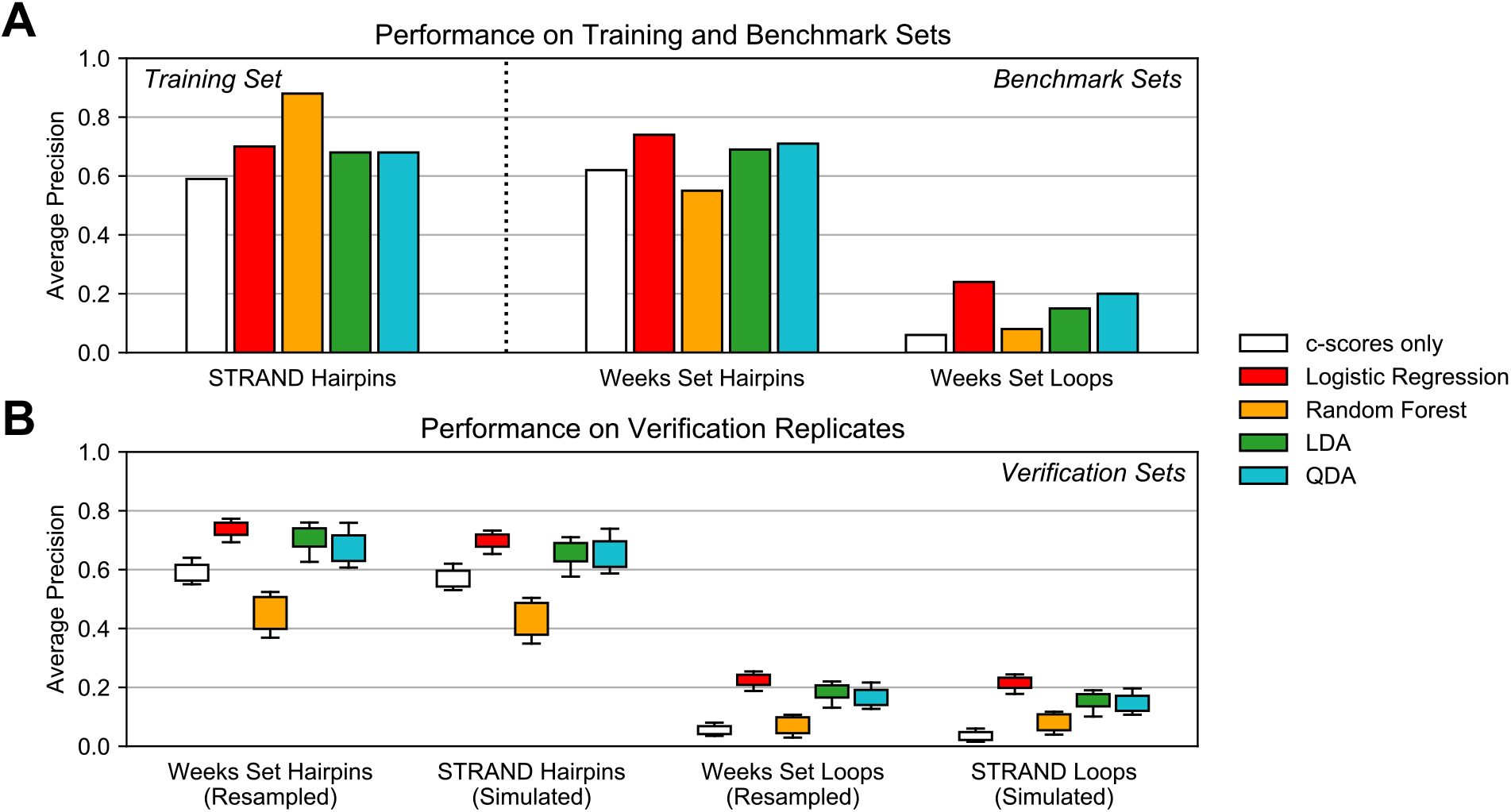
Results of experiments testing the ability of standard classifiers to fit the training set and generalize to various benchmarks and verifications. (A) Results on classifier performance on the training set (hairpins in RNA STRAND) and benchmark sets (hairpins and loops in the Weeks set). (B) Performance of trained classifiers on 5 resampled replicates of the Weeks set and 5 simulation replicates on RNA STRAND. LDA: linear discriminant analysis; QDA: quadratic discriminant analysis.

Due to these results, we decided to use an LBC trained on *c*-scores, CEL, and MEL from STRAND hairpin sites. We developed the final classifier by generating five replicates of SP data for the STRAND transcripts and using each to train a respective LBC. We benchmarked the classifiers against the Weeks set hairpins and STRAND hairpins in the other replicates and assessed their overall performance as the sum of (1) average precision on the Weeks set and (2) mean average precision on the other STRAND hairpin replicates. The classifier with the highest cumulative performance was chosen as the specific parameterization to use in *patteRNA*’s scoring, although there was little difference between the five candidates.

The final LBC performance is compiled in **Figure 4**. In short, when benchmarking on the Weeks set, we observe an increase in average precision from 0.62 with *c*-scores to 0.74, a relative improvement of almost 20%. Importantly, the precision at the highest scores (i.e., when recall is low), is significantly increased compared to *c*-scores, and roughly matches the performance seen when utilizing full transcript folding (i.e., full-length transcript partition function analysis) (see **Figure 4A**, dashed box). We confirmed that the LBC yielded slightly improved scoring when using larger contexts in computing MEL, similar to that observed in **Figure 1C**. We also utilized the entire 4,666 STRAND transcripts to benchmark *patteRNA*’s performance on various RNA classes (**Figure 4B**). As the structural properties of RNA are diverse, we observe differential performance at hairpin mining for different types of RNA. Structured transcripts defined by a high prevalence of hairpins score the best, for example, regulatory elements, small RNAs, and ribozymes. Other classes which tend to be less structured or a have large proportion of non-local base paring score relatively worse, for example, tmRNA, SRP RNA, and 5S rRNA.

**Figure 4:**
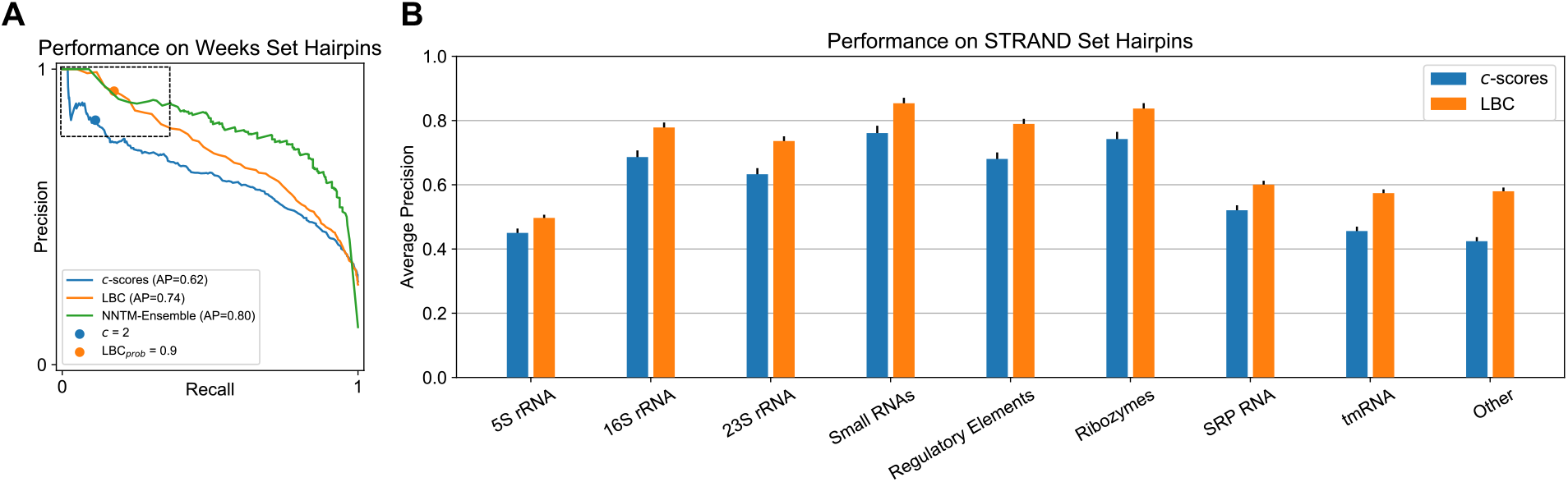
Performance of *patteRNA* when using the finalized iteration of a logistic binary classifier (LBC) natively during its scoring phase. (A) Precision-recall curves for hairpin detection in the Weeks set for LBC probabilities, full NNTM-Ensemble predictions, and regular *c*-scores absent any additional classifier processing. Dashed box indicates the region associated with the highest scores, where the LBC is able to match the precision of full-length partition function analyses. Also indicated are the performance points associated with thresholding to *c* = 2 and Prob(SL) = 0.9. (B) Average precision by RNA class when mining 5 replicated simulations on RNA STRAND transcripts. Error bars indicate standard error of the mean.

Runtime benchmarks demonstrate that our approach scales linearly and allows for transcriptome-wide mining of hairpins within an hour (see **Supplementary Figure S3**). This speed is one to two orders of magnitude faster than processing the data via local partition function workflows with windows of length 150 or 3000 nt.

### Mining Structurome Data Reveals Strong Association between Stem-Loops and RBP Binding Signals

The interplay between RNA structure and RBPs has been of significant interest for several decades (50). Such interactions are widespread, dynamic, and known to underpin important regulatory processes like splicing, trafficking, and translation (51–55). Although it is believed that many RBPs prefer to associate in unstructured regions, recent *in vitro* and *in vivo* experiments have indicated that a significant portion of RBP binding occurs in structured contexts and in a structure-dependent manner (34, 37, 56–58). That said, a mechanistic understanding of RBP binding exists only for a very small number of RBPs which have been subject to targeted research. The global trends and dynamics of RNA-protein interactions are still poorly understood, and as such, significant efforts have been directed at disentangling the complex relationships between RNA transcripts, their regulation, and the proteins which interact with them.

Corley et al. recently harnessed structure probing to detect RBP binding sites in an experiment called fSHAPE and applied it transcriptome-wide to human cell lines (34). The result of their work is a large set of data encompassing *in vivo* and *in vitro* icSHAPE reactivities and fSHAPE scores, the latter of which capture differential reactivity in the presence and absence of RBPs. They demonstrated that strong fSHAPE signals are highly correlated to RNA nucleotides that are unpaired and known to engage in hydrogen bonding with proteins, meaning that high fSHAPE signals are strong evidence of RBP binding. These data enable quantitative comparisons between RNA structure (via icSHAPE reactivity profiles) and RBP binding (high fSHAPE signals).

Corley et al.’s analysis further demonstrated that strong fSHAPE signals preferentially occur in structured contexts, and our previous work harnessing *patteRNA* expanded on this result by indicating a global association between a nucleotide-wise measure of structuredness (HDSL) and high fSHAPE (37). Both of these results, however, were obtained from “bird’s-eye view” approaches in which low-resolution global trends were utilized to elucidate general properties of RNA-protein interactions. In this work, we sought to utilize *patteRNA* to associate specific structure motifs with RBP binding in a more mechanistic “bottom-up” approach. Specifically, we sought to address the questions, “to what extent does RBP binding occur in the context of stable stem-loops?” and “what fraction of stable stem-loops can be associated with RBP binding?”

We used the LBC to score Corley et al.’s icSHAPE data as a means of exploring the association between hairpins (referred to in this section as stem-loops) and RBP binding signatures. Specifically, we mined two transcriptomes (K562 and HepG2 cells) for the representative set of stem-loops introduced earlier (stem lengths between 4 and 15 nt, loop lengths between 3 and 10 nt) and cross-referenced the locations of highly scored sites with the fSHAPE data to elucidate any connections between them. The results of our analysis are compiled in **Figure 5**, where we present findings from both *in vitro* and *in vivo* icSHAPE data (K562 results shown; results for HepG2 data were very similar and are shown in **Supplementary Figure S4**). We first examined the locations of a highly prevalent stem-loop motif described by a stem of 6 base pairs and a loop length of 4 (37). **Figure 5A** depicts the mean fSHAPE signal of highly-scoring sites (black) versus poorly-scoring sites (blue). It shows that sites which scored highly for this motif (Prob(SL) > 0.9) often display a high fSHAPE signal, interpreted as evidence of RBP binding, localized to the loop region. Specifically, greater than 70% of these high-scoring sites *in vitro* displayed strong evidence of RBP binding in the loop (defined as fSHAPE > 2, the same threshold used by Corley et al.). A score threshold of 0.9 was chosen as it is associated with near-perfect precision in our benchmarks on the Weeks set (see **Figure 4A**, orange dot). Interestingly, analysis of *in vivo* data arrives at a similar association, suggesting that data from one condition may suffice in determining relevant structures. A comparable signal was also detected when examining highly scored sites for a similar motif with a 6 base pair stem and a 3 nt loop (**Figure 5B**).

**Figure 5:**
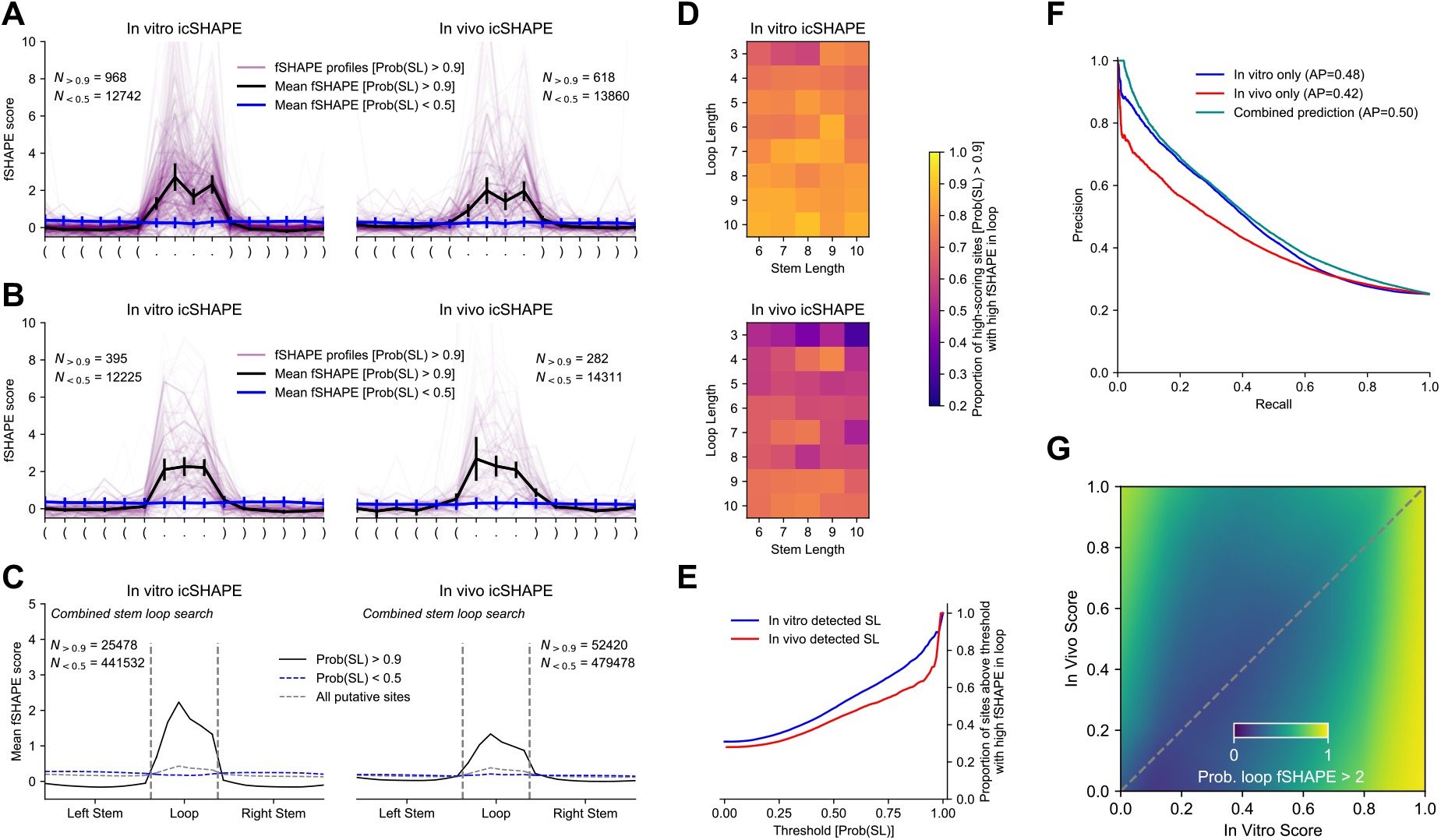
Strong association between detected stem-loops (SL) and RBP binding evidence (high fSHAPE scores) in structurome data from K562 cells. (A) fSHAPE profiles for sites scored highly (*in vitro*: left; *in vivo*: right) for a stem-loop with stem length 6 nt and loop length 4 nt. Individual fSHAPE profiles for sites with score greater than 0.9 are shown (purple) as are the mean fSHAPE profiles for sites scored above 0.9 (black) and below 0.5 (blue), respectively. (B) Same illustration as shown in panel (A), but for sites scored for a stem-loop with stem length 6 nt and loop length 3 nt. (C) Combined fSHAPE properties for sites scored when searching for a representative set of stem-loops (stem lengths 4 to 15 nt; loop lengths 3 to 10 nt; no bulges). fSHAPE profiles from scored sites were interpolated to a fixed length of 26 (10 nt left stem, 10 nt right stem, 6 nt loop; see Methods). (D) Proportion of high scoring sites (Prob(SL) > 0.9) that have fSHAPE > 2 in their predicted loop for stem-loops for each considered loop and stem length. Shown are results when mining *in vitro* icSHAPE data (top) and *in vivo* icSHAPE data (bottom). Stem-loops detected *in vitro* were more associated with evidence of RBP binding than those detected *in vivo*, but both datasets demonstrate a strong association. (E) Proportion of sites above indicated thresholds that have high fSHAPE in their predicted loop. (F) Precision-recall curves for identifying sites with high loop fSHAPE for sites that were scored in both conditions. Performance is shown for a perceptron classifier trained on condition-wise paired scores (see Methods) against the performance observed when using the condition-wise scores on their own. (G) Perceptron-modeled relationship between condition-wise scores and evidence of RBP binding (fSHAPE > 2). The modeled distribution indicates that hairpins strongly detected *in vitro* are overwhelmingly associated with RBP binding. There is also RBP binding signal identified for hairpins only detected *in vivo* and not *in vitro* (top left).

We expanded the scope of our analysis by inspecting the highly scored sites across all stem-loop motifs included in our search (motifs with stems of length 4 to 15 nt and loop length 3 to 10 nt) (see **Figure 5C**). For the *in vitro* icSHAPE data (left side of panel C), *patteRNA* identified 12,969 high scoring stem-loops out of 289,764 considered putative sites (i.e., those which satisfy sequence constraints associated with the searched motifs), which amounts to less than 5%. To visualize the fSHAPE data from these sites, which have different sizes, fSHAPE profiles were interpolated to a constant stem and loop length (10 nt and 6 nt, respectively; see Methods). When examining this larger representative set, we continued to observe a strong fSHAPE signal in loops and a low signal in stems of stable motifs. Moreover, an inspection of sites which score Prob(SL) < 0.5 shows complete depletion of this signal, thereby providing a negative control that strengthens the conclusions drawn from high-scoring sites.

Given the seemingly universal association between highly scored stem-loops and RBP binding signal, we sought to investigate it at the motif level for each considered target. In other words, we examined if particular stem-loops have a stronger association with RBP binding than others. **Figure 5D** shows the fraction of highly scored sites for each considered motif that also have high fSHAPE signal in their loop. Examining this association across all motifs reveals that this notable propensity of RBP binding signal within loops generally applies to all of them. Nevertheless, the association appears significantly stronger for *in vitro* than for *in vivo* scores. This is presumably due to the effect RBPs have on reactivities for unpaired nucleotides engaging in RBP binding (i.e., reduces their accessibility) and/or lower data quality *in vivo*. Adding to our previous conclusion that one condition may suffice for determining relevant structures (**Figure 5C**), our results indicate that *in vitro* structure mining is in fact preferable in some contexts when identifying motifs functionally relevant in an *in vivo* context. For example, differences between conditions are particularly stark when motif loops are short (e.g., top three rows of *in vivo* heatmap). We speculate that this difference is due to RBP occlusion of loop reactivities which is more detrimental to *patteRNA*’s scoring when loops are short. The differences between the conditions are further illustrated as a function of the threshold by which stem-loops are declared stable by *patteRNA* (**Figure 5E**). Notably, the observed associations were even stronger in HepG2 cells (**Supplementary Figure S4**).

Although both *in vitro* and *in vivo* detected stem-loops strongly associated with high fSHAPE signals, there were some differences between the conditions. As such, we attempted to fuse both scores into a single, more powerful predictor of stem-loops with RBP binding signals. To this end, we fitted a simple perceptron model to predict from a site’s *in vitro* and *in vivo* LBC scores whether or not the site has high fSHAPE (fSHAPE > 2) in the loop (see Methods). Using the perceptron to predict motifs with high loop fSHAPE resulted in a slightly stronger association (as quantified by average precision for indicating sites with high loop fSHAPE) between its predictions than using the *in vitro* or *in vivo* scores alone (**Figure 5F**), suggesting that changes between the two conditions can offer additional insight into RBP-motif interactions.

We attempted to interpret the perceptron’s model to gain insights into scoring patterns associated with RBP binding. Its predictions are seen in **Figure 5G** and reveal two distinct patterns. The first pattern is a high *in vitro* score (irrespective of *in vivo* score, yellow region on right side of heatmap), which recapitulates key results from **Figure 5A–E**. However, the second pattern is associated with sites that score poorly *in vitro* but strongly *in vivo* (top left corner). These sites appear to fold into stem-loops only in the *in vivo* condition. We speculate that this pattern reflects motifs that are functional (i.e., engage in RBP binding) but only fold or become stabilized in the *in vivo* cellular context. Note, however, that this pattern is far rarer than the former. Whereas over 9900 sites fall into the first pattern (Prob(SL) > 0.9 *in vitro* with high loop fSHAPE), only 66 sites were found in the second (Prob(SL) > 0.7 *in vivo*, Prob(SL) < 0.2 *in vitro*, with high loop fSHAPE). More work is needed to investigate the association of these sites with RBP binding. Note, however, that while the second pattern appeared in our analogous analysis of HepG2 data, it was not as pronounced (**Supplementary Figure S4**).

Next, we expanded the scope of our motif search to include stem-loops with bulges of 1 or 2 nt on either side of the stem. This greatly broadens the space of considered motifs and therefore increases the required computational overhead, as searching for regular stem-loops mines for 96 targets but allowing for bulges increases this number to 2,640. Overall, approximately 7.2 million sites were considered as satisfying sequence constraints for a searched motif (either a regular stem-loop or stem-loop with a bulge), 27,769 of which received a score greater than 0.9. We compiled the fSHAPE profiles of high scoring stem-loops with bulges and quantified their properties in a manner similar to our analysis on stem-loops without bulges. However, in addition to distinguishing loops from stems, we also distinguished bulges into their own group when quantifying fSHAPE tendencies. Our results are given in **Figure 6** and demonstrate a similar enrichment of high fSHAPE in apical loops of stem-loops with bulges to that which was observed for stem-loops without bulges. Moreover, we also detected a marked fSHAPE increase within bulge nucleotides, also implicating them in RBP interactions. These results expand the context of our demonstrated association between structure motifs and RBP binding signal.

**Figure 6:**
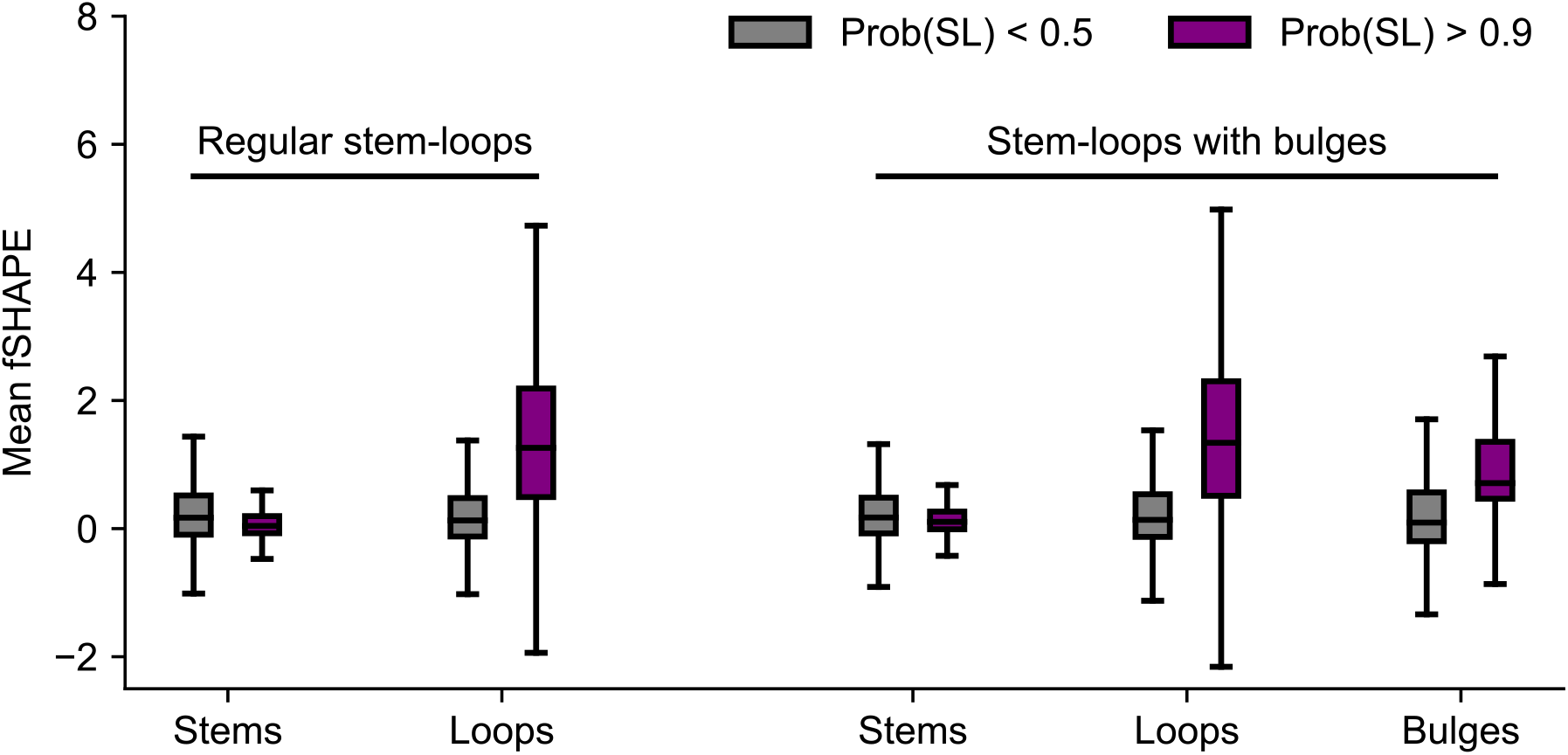
Association between RBP binding and structure motifs persists when considering stem-loop motifs with bulges in their stems. Stem-loops with bulges were defined as hairpins with stem length 4 nt to 15 nt, loop length 3 nt to 10 nt, and one bulge of 1 or 2 nt on either side of the stem.

Our analysis suggested that a significant majority of stable stem-loops likely interact with RBPs. This naturally raised the question of what fraction of RBP binding sites can be explained as occurring in the context of stem-loops. We estimated this fraction by computing the proportion of nucleotides with fSHAPE > 2 which occur in the loop segment of a highly scored SL motif in the *in vitro* data. The results are given in **Table 1**, showing that, of the fSHAPE data that was included in our motif searches, 19% of nucleotides with fSHAPE > 2 fall within a stem-loop motif scored Prob(SL) > 0.9. Upon relaxing the threshold to 0.7, this proportion increases to 33%. Interestingly, this result is comparable to previous estimates of the proportion of RBPs interacting with stem-loop motifs versus linear motifs (59). Similar results were observed *in vivo* and in HepG2 cells (see **Supplementary Table S2**) and when using NNTM-free *patteRNA c*-scores (see **Supplementary Table S3**). Nevertheless, the scope of our search remains somewhat limited. For example, we did not exhaustively consider all feasible bulge types (e.g., bulges larger than 2 nt or stem-loops with bulges on both sides of the stem), nor did we consider internal loops. Both types of motifs have been previously associated with RBPs (50, 60). Despite the computational overhead associated with mining such complicated motifs, their consideration is likely to significantly increase the proportion of high fSHAPE observations explainable as occurring in a structured element.

**Table 1:**
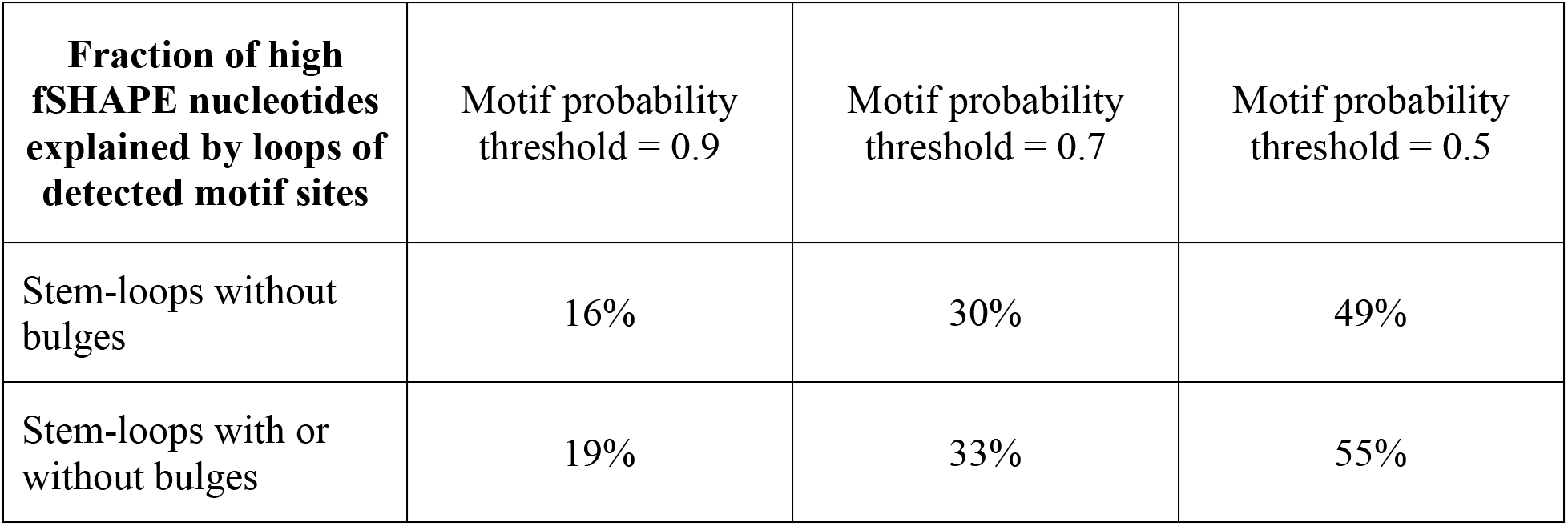
Fraction of high fSHAPE sites in K562 cells accounted for by stem-loops without bulges and stem-loops with or without bulges, as detected in *in vitro* icSHAPE data. Although the identified sites do not account for a majority of the high fSHAPE data, the results demonstrate that a sizable portion of RBP binding signals can be attributed to canonical stem-loop motifs.

Finally, we investigated the global trends of RNA-protein interactions by examining the prevalence of stem-loops within logical regions of mRNA transcripts—5’ UTRs, CDS, and 3’ UTRs (noncoding RNAs were treated as their own group). Interestingly, we observed that the RBP-SL association is remarkably consistent between regions (**Supplementary Figure S6**). Across all considered regions, approximately 75-80% of detected stem-loops had a loop which coincided with strong RBP binging signal (values indicated for K562 data; percentages were approximately 80-85% for HepG2 data). Nevertheless, we observed large differences in the density of detected stem-loops between these regions. In all cell lines and conditions, 3’ UTRs have a significantly higher rate of stable stem-loops than other regions (see **Table 2**). For instance, in K562 *in vivo* icSHAPE data, *patteRNA* identified 9.57 stem-loops per 1000 nt in 3’ UTRs, compared to 4.86 and 4.07 in 5’ UTRs and CDS, respectively. In the context of post-transcriptional regulation, stem-loops are known to be mechanistically involved with polyadenylation and degradation (61–63); however, this is the first stem-loop profile of a human structurome that systematically quantifies this at a global level. In the context of our observations that the association between stem-loops and RBP binding is fairly uniform across logical transcript regions and there is an increased prevalence of stem-loops in 3’ UTRs, one can infer that stem-loops and RBP are globally involved in 3’ UTR regulatory processes like polyadenylation and degradation. Our finding warrants the continued investigation of local structure motifs in 3’ UTRs and their roles in the interactome.

**Table 2:** Density of stem-loop detections in logical regions of mRNA transcripts from *in vitro* and *in vivo* icSHAPE data. Values are given as stem-loops per 1000 nt.

|  | Region | K562 | HepG2 |
| --- | --- | --- | --- |
| *In vitro* icSHAPE | 5' UTRs | 7.33 | 4.84 |
|  | CDS | 8.71 | 5.71 |
|  | 3' UTRs | 11.63 | 8.61 |
|  | ncRNAs | 9.78 | 6.31 |
| *In vivo* icSHAPE | 5' UTRs | 4.86 | 4.52 |
|  | CDS | 4.07 | 3.99 |
|  | 3' UTRs | 9.57 | 6.98 |
|  | ncRNAs | 6.74 | 4.71 |

## DISCUSSION

The evolution and growing scale of RNA structure probing experiments has warranted methods well-suited to the analysis of millions to billions of nucleotides. *patteRNA* is one such tool which was developed with the specific aim of rapidly extracting biologically relevant insights from such data. For genome-wide analyses, high precision is often a specific objective yet challenging to achieve due to the large number of negative sites considered (64). In this work, we took a machine learning approach to improve scoring precision by developing a classifier that accounts for local sequence energetics in addition to *patteRNA*’s statistical characterization of reactivities. To ensure broad applicability, we created a high-quality, non-redundant, and large-scale set of transcripts with known structures from RNA STRAND and used it in conjunction with a data simulation scheme to extensively train and validate our approach. Our work indicates this simulated data provide a strong suite of structural information by which to develop methods, which can augment real datasets that are currently much smaller in size. We believe this resource will be useful for others seeking to develop data-driven methods. Application of the classifier transcriptome-wide revealed that stable stem-loops are strongly associated with fSHAPE RBP binding signals across cell lines. This association has been previously documented for individual RBPs (56, 59, 60), however the ubiquitous nature of stem-loops to interact with RBPs *in vivo* has not been previously shown. Not only does this implicate common and canonical structural elements with RBPs, it also reinforces the notion that mining local structure elements can provide biologically relevant insights.

The results of our perceptron analysis of condition-wise paired scores demonstrated that some patterns could be leveraged to identify functional stem-loops beyond inspecting each condition independently. We found that stem-loops detected *in vitro* explain a significant (greater than 30%) fraction of RBP binding signals in Corley et al.’s data. Another pattern that emerged was the presence of stem-loops that score poorly *in vitro* (Prob(SL) < 0.2), but highly *in vivo* (Prob(SL) > 0.7), although the prevalence of such sites was much lower than sites associated with the former pattern. We note that the perceptron analysis was primarily performed to assist in the interpretation of score changes between conditions (e.g., **Figure 5G**), and that analogous statistical analysis (i.e., via bivariate data fitting) could arrive at similar conclusions. Lastly, it is possible that a more advanced perceptron approach could better disentangle the relationship between the two conditions. For instance, a perceptron or deep neural network trained on the underlying features from each site in each condition (i.e., *c*-score, MEL, CEL, etc.) might yield more precise predictions on the identification of structure motifs with RBP binding signal.

The LBC developed in this work was demonstrated to be significantly more accurate than *patteRNA*’s *c*-scores alone. Nevertheless, we were curious to what degree using *c*-scores (i.e., an NNTM-free approach) could recapitulate the stem-loop/RBP results obtained with the LBC. We re-analyzed the data, but used a threshold of *c* > 2 to determine stable stem-loops instead of Prob(SL) > 0.9 (see **Supplementary Figure S5**). As indicated on **Figure 4A**, this threshold is roughly comparable to an LBC threshold of 0.9, although it yields slightly lower precision and recall. We found that the use of *c*-scores arrived at similar conclusions to those which were obtained with the LBC, but the observed association was slightly weaker. Specifically, we observed that the association was significantly weaker for stem-loops with shorter stems (6 or 7 nt) and longer loops (5 nt or longer), especially for the *in vivo* data (**Supplementary Figure S5F**). We believe that such motifs benefit most from the thermodynamic information contained in MEL, as sequence constraints are less effective in pruning the number of negative sites considered during scoring. Nevertheless, this result recapitulates that *patteRNA*’s NNTM-free implementation provides accurate detections, especially for high-quality data. We believe that the LBC assists most in situations where motifs are short, or data quality is low.

*patteRNA* was developed with a specific aim of addressing the need for universal and efficient tools for analysis of a growing breadth, scale, and diversity of SP experiments. Universality is important because different experiments yield reactivities with disparate statistical properties, meaning one-size-fits-all approaches are generally suboptimal. As such, the versatility of *patteRNA* is a central characteristic of the method. In the development of a data-driven scoring approach, we sought to maintain this trait. We found that the *c*-score naturally lends itself to ensuring an automatically adaptable classifier, as it provides a normalized measure of a site’s consistency with the target motif. Serving as a measure of statistical significance against a null distribution that captures data-level and motif-level biases, this metric can be considered largely data-invariant. Our previous work further demonstrated the suitability of *patteRNA*’s comparative scores to accurately characterize the presence structure of motifs in a diverse suite of simulated reactivity models (37). Moreover, the MEL feature only depends on the local nucleotide sequence, meaning that it is invariant to different reactivity distributions. The decoupling of MEL from the SP data also enables insight on the contributions of NNTM to SP data interpretation. For example, although we observed the LBC improves precision across a range of motifs, the largest relative improvement was observed for motifs with few base pairs, such as loops flanked by single base pair (see **Figure 3**). This trend was also observed in our analysis of the Corley data, where the largest differences between using *c*-scores (NNTM-free) and the LBC (NNTM-dependent) was observed for the shortest stems. Our results suggest that, when searching for motifs harboring many base pairs, folding with NNTM may not provide a significant benefit over using *c*-scores alone.

From its initial development, *patteRNA* was not envisioned as a replacement or competing method to traditional NNTM-based approaches typically used in RNA structure analyses. Rather, it was developed as a tool to be used in tandem to NNTM-based approaches. For example, it can be used to identify candidate sites for a motif of interest (e.g., broadly defined motifs, such as stem-loops, or specific structural elements, such as iron response elements), which could then be subject to more intensive structural analysis with NNTM and targeted SP experiments. In any case, the advantages of *patteRNA* emerge when analyzing large-scale data. By focusing specifically on sites that satisfy the sequence constraints for a target motif and performing minimal local MFE calculations for the LBC, our method arrives at structuromic insights orders of magnitude faster than partition-function based analyses. This speed helps mitigate the computational overhead associated with partition-function analysis of massive SP datasets, especially for those without access to cutting-edge computational hardware. In considering the future development of a method like *patteRNA*, we believe more work remains to be done, despite *patteRNA*’s demonstrated capabilities and scalability relative to transcript folding or partition function analysis. The primary limitation of our method is the dependence on the definition of specific local secondary structure motifs to use for mining. This dependence enables rapid scans in large datasets but limits the scope of the method’s analysis to elements with a previously known or suspected structure. One may specify a large set of related motifs to circumvent this limitation, but this comes at an increased computational cost. The current implementation is capable of mining thousands of distinct structures in a human transcriptome within several hours (e.g., mining stem-loops with bulges), however searches with increased flexibility (e.g., accounting for more diverse bulges, longer loops, and internal loops) result in a combinatorial explosion of considered motifs to counts larger than 10,000 or 100,000. This renders such searches impractical. Nevertheless, when specifically focused on canonical local motifs, for example stem-loops or stem-loops with bulges, *patteRNA* provides rapid, accurate, and biologically relevant motif mining capabilities on structurome data.

NNTM-based methods have recently seen the application of pattern recognition in SHAPE data to improve the quality of secondary structure prediction. SHAPELoop (65) generates a Boltzmann ensemble and selects suboptimal structures in which loops are more consistent with SHAPE data than the MFE based on a learned set of loop reactivity patterns. Their approach, similarly to *patteRNA*, integrates single nucleotide observations to the motif level, which resulted in structure predictions with a slightly higher accuracy than standard approaches for data-directed folding. Although their pattern models were extracted from a small set of reference structures, they were shown to correlate with stereochemical properties of SHAPE, suggesting broader applicability to other SP probes. Nevertheless, their patterns are specific to SHAPE and likely require recalibration for different probing methodologies. This relies on the availability of reference structures with high-quality data, which are currently lacking. In the context of a tool like *patteRNA*, SHAPELoop utilizes SP data towards an entirely different objective (structure prediction *de novo* versus structure mining), but the results of their work demonstrate that stereochemically-correlated reactivity information can be utilized to more accurately identify loops. The utilization of such information when scanning large SP data for motifs containing loops would likely improve *patteRNA*’s accuracy. The pattern models of SHAPELoop, derived from a statistical analysis of reference data, should be naturally integrable to the data-driven approach developed in this work; namely, the LBC naturally extends to additional features capturing, for example, loop-specific pattern scores. The previous design of *patteRNA* (37), however, would render such integration non-trivial. Nevertheless, the incorporation of such stereochemical trends in an automated and universal way (i.e., generalization outside of *in vitro* SHAPE data) is an unaddressed challenge. Moreover, identification of specific stereochemical patterns could likely assist in furthering our understanding of biological structures, such as the precise structural context underpinning interactions like RNA-protein binding.

## Data Availability

The latest version of *patteRNA*, version 2.1, was used for all analyses in this study. *patteRNA* is an open-source Python 3 module and is freely available at www.github.com/AviranLab/patteRNA under the BSD-2 license. The data that support the findings of this study are openly available in Zenodo at http://doi.org/10.5281/zenodo.4667910, reference number (44).

## Competing Interests

No potential competing interest was reported by the authors.

## Notes

### Competing Interest Statement

The authors have declared no competing interest.

## REFERENCES

1. Ganser,L.R., Kelly,M.L., Herschlag,D. and Al-Hashimi,H.M. (2019) The roles of structural dynamics in the cellular functions of RNAs. Nat. Rev. Mol. Cell Biol., 20, 474–489.

2. Mustoe,A.M., Brooks,C.L. and Al-Hashimi,H.M. (2014) Hierarchy of RNA functional dynamics. Annu. Rev. Biochem., 83, 441–466.

3. Fica,S.M. and Nagai,K. (2017) Cryo-electron microscopy snapshots of the spliceosome: Structural insights into a dynamic ribonucleoprotein machine. Nat. Struct. Mol. Biol., 24, 791–799.

4. Dallaire,P., Tan,H., Szulwach,K., Ma,C., Jin,P. and Major,F. (2016) Structural dynamics control the MicroRNA maturation pathway. Nucleic Acids Res., 44, 9956–9964.

5. Serganov,A. and Patel,D.J. (2007) Ribozymes, riboswitches and beyond: Regulation of gene expression without proteins. Nat. Rev. Genet., 8, 776–790.

6. Esteller,M. (2011) Non-coding RNAs in human disease. Nat. Rev. Genet., 12, 861–874.

7. Battle,D.J. and Doudna,J.A. (2001) The stem-loop binding protein forms a highly stable and specific complex with the 3’ stem-loop of histone mRNAs. Rna, 7, 123–132.

8. Šponer,J., Bussi,G., Krepl,M., Banáš,P., Bottaro,S., Cunha,R.A., Gil-Ley,A., Pinamonti,G., Poblete,S., Jurečka,P., et al. (2018) RNA Structural Dynamics As Captured by Molecular Simulations: A Comprehensive Overview. Chem. Rev., 118, 4177–4338.

9. Holbrook,S.R. and Kim,S.-H. (1997) RNA crystallography. Biopolymers, 44, 3–21.

10. Fürtig,B., Richter,C., Wöhnert,J. and Schwalbe,H. (2003) NMR Spectroscopy of RNA. ChemBioChem, 4, 936–962.

11. Zhang,K., Li,S., Kappel,K., Pintilie,G., Su,Z., Mou,T.-C., Schmid,M.F., Das,R. and Chiu,W. (2019) Cryo-EM structure of a 40 kDa SAM-IV riboswitch RNA at 3.7 Å resolution. Nat. Commun., 10, 5511.

12. Pace,N.R., Thomas,B.C. and Woese,C.R. (1999) Probing RNA structure, function, and history by comparative analysis. Cold Spring Harb. Monogr. Ser., 37, 113–142.

13. Gutell,R.R., Lee,J.C. and Cannone,J.J. (2002) The accuracy of ribosomal RNA comparative structure models. Curr. Opin. Struct. Biol., 12, 301–310.

14. Larsen,N. and Zwieb,C. (1991) SRP-RNA sequence alignment and secondary structure. Nucleic Acids Res., 19, 209–215.

15. Nussinov,R. and Jacobson,A.B. (1980) Fast algorithm for predicting the secondary structure of single-stranded RNA. Proc. Natl. Acad. Sci. U. S. A., 77, 6309–6313.

16. Zuker,M. (1989) On finding all suboptimal foldings of an RNA molecule. Science (80-.)., 244, 48–52.

17. Gardner,P.P. and Giegerich,R. (2004) A comprehensive comparison of comparative RNA structure prediction approaches. BMC Bioinformatics, 5, 140.

18. Knapp,G. (1989) Enzymatic approaches to probing of RNA secondary and tertiary structure. In Methods in Enzymology. Academic Press, Vol. 180, pp. 192–212.

19. Wilkinson,K.A., Merino,E.J. and Weeks,K.M. (2006) Selective 2’-hydroxyl acylation analyzed by primer extension (SHAPE): quantitative RNA structure analysis at single nucleotide resolution. Nat. Protoc., 1, 1610–1616.

20. Choudhary,K., Deng,F. and Aviran,S. (2017) Comparative and integrative analysis of RNA structural profiling data: current practices and emerging questions. Quant. Biol., 5, 3–24.

21. Weng,X., Gong,J., Chen,Y., Wu,T., Wang,F., Yang,S., Yuan,Y., Luo,G., Chen,K., Hu,L., et al. (2020) Keth-seq for transcriptome-wide RNA structure mapping. Nat. Chem. Biol., 16, 489–492.

22. Wang,P.Y., Sexton,A.N., Culligan,W.J. and Simon,M.D. (2019) Carbodiimide reagents for the chemical probing of RNA structure in cells. RNA, 25, 135–146.

23. Marinus,T., Fessler,A.B., Ogle,C.A. and Incarnato,D. (2021) A novel SHAPE reagent enables the analysis of RNA structure in living cells with unprecedented accuracy. Nucleic Acids Res., 10.1093/nar/gkaa1255.

24. Aviran,S., Trapnell,C., Lucks,J.B., Mortimer,S.A., Luo,S., Schroth,G.P., Doudna,J.A., Arkin,A.P. and Pachter,L. (2011) Modeling and automation of sequencing-based characterization of RNA structure. Proc. Natl. Acad. Sci. U. S. A., 108, 11069–11074.

25. Deigan,K.E., Li,T.W., Mathews,D.H. and Weeks,K.M. (2009) Accurate SHAPE-directed RNA structure determination. Proc. Natl. Acad. Sci. U. S. A., 106, 97–102.

26. Lorenz,R., Luntzer,D., Hofacker,I.L., Stadler,P.F. and Wolfinger,M.T. (2016) SHAPE directed RNA folding. Bioinformatics, 32, 145–147.

27. Mustoe,A.M., Busan,S., Rice,G.M., Hajdin,C.E., Peterson,B.K., Ruda,V.M., Kubica,N., Nutiu,R., Baryza,J.L. and Weeks,K.M. (2018) Pervasive Regulatory Functions of mRNA Structure Revealed by High-Resolution SHAPE Probing. Cell, 173, 181–195.e18.

28. Twittenhoff,C., Brandenburg,V.B., Righetti,F., Nuss,A.M., Mosig,A., Dersch,P. and Narberhaus,F. (2020) Lead-seq: Transcriptome-wide structure probing in vivo using lead(II) ions. Nucleic Acids Res., 48, E71–E71.

29. Choudhary,K., Lai,Y.-H., Tran,E.J. and Aviran,S. (2019) dStruct: identifying differentially reactive regions from RNA structurome profiling data. Genome Biol., 20, 40.

30. Marangio,P., Law,K.Y.T., Sanguinetti,G. and Granneman,S. (2020) Differential BUM-HMM: a robust statistical modelling approach for detecting RNA flexibility changes in high-throughput structure probing data. bioRxiv, 10.1101/2020.07.30.229179.

31. Saha,K., England,W., Fernandez,M.M., Biswas,T., Spitale,R.C. and Ghosh,G. (2020) Structural disruption of exonic stem-loops immediately upstream of the intron regulates mammalian splicing. Nucleic Acids Res., 48, 6294–6309.

32. Yang,M., Woolfenden,H.C., Zhang,Y., Fang,X., Liu,Q., Vigh,M.L., Cheema,J., Yang,X., Norris,M., Yu,S., et al. (2020) Intact RNA structurome reveals mRNA structure-mediated regulation of miRNA cleavage in vivo. Nucleic Acids Res., 48, 8767–8781.

33. Ding,Y., Tang,Y., Kwok,C.K., Zhang,Y., Bevilacqua,P.C. and Assmann,S.M. (2014) In vivo genome-wide profiling of RNA secondary structure reveals novel regulatory features. Nature, 505, 696–700.

34. Corley,M., Flynn,R.A., Lee,B., Blue,S.M., Chang,H.Y. and Yeo,G.W. (2020) Footprinting SHAPE-eCLIP Reveals Transcriptome-wide Hydrogen Bonds at RNA-Protein Interfaces. Mol. Cell, 80, 903–914.e8.

35. Ledda,M. and Aviran,S. (2018) PATTERNA: Transcriptome-wide search for functional RNA elements via structural data signatures. Genome Biol., 19, 28.

36. Radecki,P., Ledda,M. and Aviran,S. (2018) Automated recognition of RNA structure motifs by their SHAPE data signatures. Genes (Basel)., 9, 300.

37. Radecki,P., Uppuluri,R. and Aviran,S. (2021) Rapid structure-function insights via hairpin-centric analysis of big RNA structure probing datasets. bioRxiv, 10.1101/2021.04.27.441661.

38. Deng,F., Ledda,M., Vaziri,S. and Aviran,S. (2016) Data-directed RNA secondary structure prediction using probabilistic modeling. RNA, 22, 1109–1119.

39. Andronescu,M., Bereg,V., Hoos,H.H. and Condon,A. (2008) RNA STRAND: The RNA secondary structure and statistical analysis database. BMC Bioinformatics, 9, 340.

40. Hajdin,C.E., Bellaousov,S., Huggins,W., Leonard,C.W., Mathews,D.H. and Weeks,K.M. (2013) Accurate SHAPE-directed RNA secondary structure modeling, including pseudoknots. Proc. Natl. Acad. Sci. U. S. A., 110, 5498–5503.

41. Lavender,C.A., Lorenz,R., Zhang,G., Tamayo,R., Hofacker,I.L. and Weeks,K.M. (2015) Model-Free RNA Sequence and Structure Alignment Informed by SHAPE Probing Reveals a Conserved Alternate Secondary Structure for 16S rRNA. PLOS Comput. Biol., 11, e1004126.

42. Huang,Y., Niu,B., Gao,Y., Fu,L. and Li,W. (2010) CD-HIT Suite: a web server for clustering and comparing biological sequences. Bioinformatics, 26, 680–682.

43. Sükösd,Z., Swenson,M.S., Kjems,J. and Heitsch,C.E. (2013) Evaluating the accuracy of SHAPE-directed RNA secondary structure predictions. Nucleic Acids Res., 41, 2807–2816.

44. Radecki,P., Uppuluri,R., Deshpande,K. and Aviran,S. (2021) Dataset: Accurate Detection of RNA Stem-Loops in Structurome Data Reveals Widespread Association with Protein Binding Sites. 10.5281/zenodo.4667910, 10.5281/ZENODO.4667910.

45. Kingma,D.P. and Ba,J. (2014) Adam: A Method for Stochastic Optimization. arXiv.

46. Rabiner,L.R. (1989) A tutorial on hidden Markov models and selected applications in speech recognition. Proc. IEEE, 77, 257–286.

47. Rouskin,S., Zubradt,M., Washietl,S., Kellis,M. and Weissman,J.S. (2014) Genome-wide probing of RNA structure reveals active unfolding of mRNA structures in vivo. Nature, 505, 701–705.

48. Li,H. and Aviran,S. (2018) Statistical modeling of RNA structure profiling experiments enables parsimonious reconstruction of structure landscapes. Nat. Commun., 9, 606.

49. Ding,Y. and Lawrence,C.E. (2003) A statistical sampling algorithm for RNA secondary structure prediction. Nucleic Acids Res., 31, 7280–7301.

50. Corley,M., Burns,M.C. and Yeo,G.W. (2020) How RNA-Binding Proteins Interact with RNA: Molecules and Mechanisms. Mol. Cell, 78, 9–29.

51. Sysoev,V.O., Fischer,B., Frese,C.K., Gupta,I., Krijgsveld,J., Hentze,M.W., Castello,A. and Ephrussi,A. (2016) Global changes of the RNA-bound proteome during the maternal-to-zygotic transition in Drosophila. Nat. Commun., 7, 12128.

52. Gebauer,F., Schwarzl,T., Valcárcel,J. and Hentze,M.W. (2021) RNA-binding proteins in human genetic disease. Nat. Rev. Genet., 22, 185–198.

53. Wei,C., Xiao,R., Chen,L., Cui,H., Zhou,Y., Xue,Y., Hu,J., Zhou,B., Tsutsui,T., Qiu,J., et al. (2016) RBFox2 Binds Nascent RNA to Globally Regulate Polycomb Complex 2 Targeting in Mammalian Genomes. Mol. Cell, 62, 875–889.

54. Castello,A., Fischer,B., Eichelbaum,K., Horos,R., Beckmann,B.M., Strein,C., Davey,N.E., Humphreys,D.T., Preiss,T., Steinmetz,L.M., et al. (2012) Insights into RNA Biology from an Atlas of Mammalian mRNA-Binding Proteins. Cell, 149, 1393–1406.

55. Gerstberger,S., Hafner,M. and Tuschl,T. (2014) A census of human RNA-binding proteins. Nat. Rev. Genet., 15, 829–845.

56. Dominguez,D., Freese,P., Alexis,M.S., Su,A., Hochman,M., Palden,T., Bazile,C., Lambert,N.J., Nostrand,E.L. Van, Pratt,G.A., et al. (2018) Sequence, Structure, and Context Preferences of Human RNA Binding Proteins. Mol. Cell, 70, 854–867.e9.

57. Nostrand,E.L. Van, Freese,P., Pratt,G.A., Wang,X., Wei,X., Xiao,R., Blue,S.M., Chen,J.-Y., Cody,N.A.L., Dominguez,D., et al. (2020) A large-scale binding and functional map of human RNA-binding proteins. Nature, 583, 711–719.

58. Sun,L., Xu,K., Huang,W., Yang,Y.T., Li,P., Tang,L., Xiong,T. and Zhang,Q.C. (2021) Predicting dynamic cellular protein–RNA interactions by deep learning using in vivo RNA structures. Cell Res., 10.1038/s41422-021-00476-y PMID - 33623109.

59. Jolma,A., Zhang,J., Mondragón,E., Morgunova,E., Kivioja,T., Laverty,K.U., Yin,Y., Zhu,F., Bourenkov,G., Morris,Q., et al. (2020) Binding specificities of human RNA-binding proteins toward structured and linear RNA sequences. Genome Res., 30, 962–973.

60. Maticzka,D., Lange,S.J., Costa,F. and Backofen,R. (2014) GraphProt: modeling binding preferences of RNA-binding proteins. Genome Biol., 15, R17.

61. Kuhn,J., Tengler,U. and Binder,S. (2001) Transcript Lifetime Is Balanced between Stabilizing Stem-Loop Structures and Degradation-Promoting Polyadenylation in Plant Mitochondria. Mol. Cell. Biol., 21, 731–742.

62. Phillips,C., Kyriakopoulou,C.B. and Virtanen,A. (1999) Identification of a stem-loop structure important for polyadenylation at the murine IgM secretory poly(A) site. Nucleic Acids Res., 27, 429–438.

63. Grechishnikova,D. and Poptsova,M. (2016) Conserved 3’ UTR stem-loop structure in L1 and Alu transposons in human genome: possible role in retrotransposition. BMC Genomics, 17, 992.

64. Eddy,S.R. (2014) Computational analysis of conserved RNA secondary structure in transcriptomes and genomes. Annu. Rev. Biophys., 43, 433–456.

65. Cao,J. and Xue,Y. (2021) Characteristic chemical probing patterns of loop motifs improve prediction accuracy of RNA secondary structures. Nucleic Acids Res., 10.1093/nar/gkab250 PMID - 33849076.

